# SABRE: Self-Attention Based model for predicting T-cell Receptor Epitope Specificity

**DOI:** 10.1101/2023.10.02.560555

**Authors:** Zicheng Wang, Yufeng Shen

## Abstract

T cell receptors (TCR) recognize antigens on the surface of T cells, which is the critical event in the adaptive immune response to infection and vaccination. The ability to determine TCR-antigen recognition would benefit research in basic immunology and therapeutics. High-throughput experimental approaches for determining TCR-antigen specificity have produced valuable data, but the TCR-antigen pairing space is astronomically more significant than what can reached by experiments. Here, we describe a computational method for predicting TCR-antigen recognition, SABRE (Self-Attention-based Transformer Model for predicting T-cell Receptor-Epitope specificity). SABRE captures sequence properties of matching TCR and antigen pairs by selfsupervised pre-training using known pairs from curated databases and large-scale experiments. It then fine-tunes by supervised learning to predict TCRs that can recognize each antigen. We showed that SABRE’s AUROC reaches 0.726 ± 0.008 for predicting TCR-epitope recognition. We meticulously designed a training and testing scheme to evaluate the model’s performance on unseen TCR species: 60% of the data was allocated for training, 20% for validation, and the remaining 20% exclusively for testing. Notably, this testing set comprised entirely of TCRs not present in the training phase, ensuring a genuine assessment of the model’s ability to generalize to novel data.

## Introduction

The binding of peptide-MHC complex by T cell receptors is crucial in T cell activation during adaptive immune responses ^1,2^. Predicting TCR-peptide binding is crucial due to its potential in various health applications, like cancer immunotherapies ^3-5^, vaccine design ^6,7^, and autoimmune disease understanding ^8-10^. Additionally, it enhances public health strategies by predicting immune responses ^11,12^ and supports personalized medicine ^13,14^. Despite its complexity, computational biology and machine learning advancements make accurate predictions increasingly feasible.

T cell receptor for epitope binds both peptide and MHC determinants. Consequently, these receptors’ antigenic peptides and specificity are often presented as unique in the immune system ^15,16^. Furthermore, the thymus’s positive and negative selection processes result in diverse T cells in the naive repertoire that can recognize such peptide-MHC complexes ^17^. Though we know the binding specificity is determined by each part of the complex (TRA, TRB, peptide, and MHC), the contribution is different. Beta CDR3, the V gene and J gene correspond to the leading contributors, followed by the alpha chain CDR3 and MHC, and therefore no contribution to the cell type ^18^. Thus, well measurement of TCR and epitope binding mainly focuses on the beta CDR3.

Though CDR3 and epitope are short, the potential number of the sequences is huge. There are effectively 13^20 - 14^20 possible CDR3 sequences, and the key issue is that we can only sample a tiny fraction of TCRs in any experiments. This makes using prediction ways to model the interaction important. Computational tools for analyzing TCR patterns and predicting peptide–TCR binding fall into three categories. The first, including tools like TCRdist ^19^, GLIPH ^20^, and DeepTCR ^21^, use clustering to uncover antigen-specific binding patterns but doesn’t directly identify peptide– TCR binding. The second, featuring models like TCRGP ^22^, and TCRex ^23^ predict TCR binding to specific peptides, limiting their applicability. The third group, with tools such as ERGO2 ^24^, pMTnet ^25^, TITAN ^26^, and PanPep ^27^ build broader prediction models but struggle with specific identification questions. To create an easily finetuned and quickly applied model for different epitopes, In this study, we developed a prediction model based on self-attention models ^28-30^ to learn a representation of TCR and epitope sequences based on VDJdb ^31^ and MIRA databases ^32^. Following the pre-trained transformer encoder, we added classification layers to predict the specificity of unseen TCRs for known epitopes. Moreover, we created several epitope-specific models using finetuning techniques ^33,34^. This task involved retraining and testing the model for a particular epitope. We demonstrated that the flexible and fast finetuned model could be easily used for specific epitope-TCR for precision prediction.

## Result

We developed SABRE, a Self-Attention Based model for predicting T-cell Receptor Epitope Specificity. The model has two components: a pre-trained transformer encoder and a binary classifier. To establish the model, data are from two sources: the VDJdb, a general TCR-epitope database, and the MIRA database, which is specific to COVID-19. After we processed the TCR-epitope, we divided the data into three sets: the training. Test and validation datasets. None of the TCRs was shared among the three datasets, and the pretraining part adopted training and validation datasets as input, while the classification part distinguished training and validation datasets for different purposes. Test TCRs were absent in the pretraining part. In classification, each epitope in three datasets was combined with five random TCRs from collected individuals’ repertoires to form the opposing control pairs. Training, validation, and test data split for each source fellow the ratio 3:1:1. Single TRB in the MIRA dataset could react to multiple epitopes; therefore, the unique number of TCR was less than that of TRB-epitope pairs.

The TRB CDR3 sequence was paired with the epitope input, tokenized by amino acid, and concatenated with a unique character. The intuition behind using a transformer for pre-training is to capture intricate relationships and learn a robust representation of key residues and their contextual importance in TCR-antigen pairs. The pre-training procedure uses a transformer model to predict masked positions from sequence encoding based on the multi-attention mechanism. We concatenate the paired TCR CDR3 and epitope sequences as the input. SABRE employed a highdimensional vector to represent CDR3 and epitope sequences encoded by the transformer module. By incorporating a classification deep neural network atop the encoder, the model was fine-tuned for prediction. The SABRE model attained an AUC of 0.726 ± 0.008, outperforming the benchmark distance-based method (comparing the edit distance between training and test TCR, regarding them binding to the same epitope if the distance is less or equal to one, see Method), which achieved 0.580 ± 0.006. The distance-based method exhibited a low TPR (0.00149) despite reasonably controlling false positive rates (0) when employing a distance threshold of one. We compared the SABRE model to state-of-the-art methods such as PepPan. After 1,000 fine-tuning steps, the AUROC for the majority setting of PanPep in predicting the same dataset was 0.703.

In contrast, fewer or more steps resulted in lower performance (AUROC of 0.690 and 0.676, respectively). The few-shot setting yielded an AUROC of 0.642. Additionally, we utilized TITAN’s robust baseline, the k-nearest-neighbor (K-NN) classifier, as a supplementary benchmark, which achieved an AUC no more significant than 0.56.

The accurate prediction of TCR-epitope interactions was constrained by the availability of training data covering the entire sequence space. To enhance our model’s performance, we employed a fine-tuning strategy. We first trained a model on a comprehensive dataset capturing diverse TCR-epitope interactions. Then, we further refined this pre-trained model to create epitope-specific versions. We selected several high-abundance epitopes (at least 200 related unique TCRs) to tackle this challenge and fine-tuned the pre-trained model for each. Our results indicated that prediction accuracy varies significantly among epitopes. While the prediction performance for specific epitopes’ TCRs achieved an AUROC of 0.9 (e.g., YLQ from SARS-CoV-2 spike protein), others were impossible to predict accurately. This variation made it difficult to surpass an overall prediction accuracy of 0.75 or higher. We recommend carefully considering the data sources required for hard-to-predict TCRs associated with specific epitopes to improve performance.

We compared the performance of general and specific models across 18 large recorders of epitopes from the VDJdb. Most epitope-specific models outperformed the general one, especially for the KAF from the HIV-1 Gag gene, where the specific model improved AUROC by more than 30%. However, epitope-specific models may have encountered overfitting issues. For IVT (EBV EBNA4), the specific model did not outperform the general one. Furthermore, for some epitopes like ELA (Homo sapiens MLANA), LLW (YFV NS4B), and TTD (SARS-CoV-2 NSP3), their specific models slightly underperformed compared to the general models.

Several factors might cause the varying performance of each epitope, regardless of specific or general models. We found no direct evidence that the number of training sizes impacts the result. The number of related TCRs of each epitope was not correlated with the AUC (Pearson correlation equals -0.0247, p-value equals 0.922, Supplementary Figure 5). We also demonstrated that TCR similarity between training and test data highly correlated with performance. Smaller distances (measured using edit distance) improved performance (Pearson correlation coefficient equals - 0.817, p-value equals 3.6*10-5).

To fine-tune the pretraining model on the MIRA dataset, we obtained more than 150,000 high-confidence SARS-CoV-2-specific TCR-epitope pairs. However, most TCRs recognize multiple epitopes, and epitopes are distributed across different areas of the virus genome, with TCRs originating from various donors. The MIRA dataset consists of 269 epitope pools and 545 unique epitopes. We used epitopes with many TCRs (at least 200 pairs) to fine-tune the COVID epitope-specific model. Compared with the general model prediction (downsample 200 pairs for each epitope to train the general model), most specific models (using 60% of all pairs for specific epitopes to train the specific model) outperform the general one (188 out of 287). Especially for the epitope high abundance models (at least 2000 pairs), 55 out of 59 specific models predicting AUCs are better than the general model. However, the highest number of epitope pairs didn’t mean the best performance. IEL/LID/IDF specific models (the number of pairs is over 17,000) predicted AUC was around 0.72, which stood in the middle across all specific models (Figure 3a).

**Figure 1.**
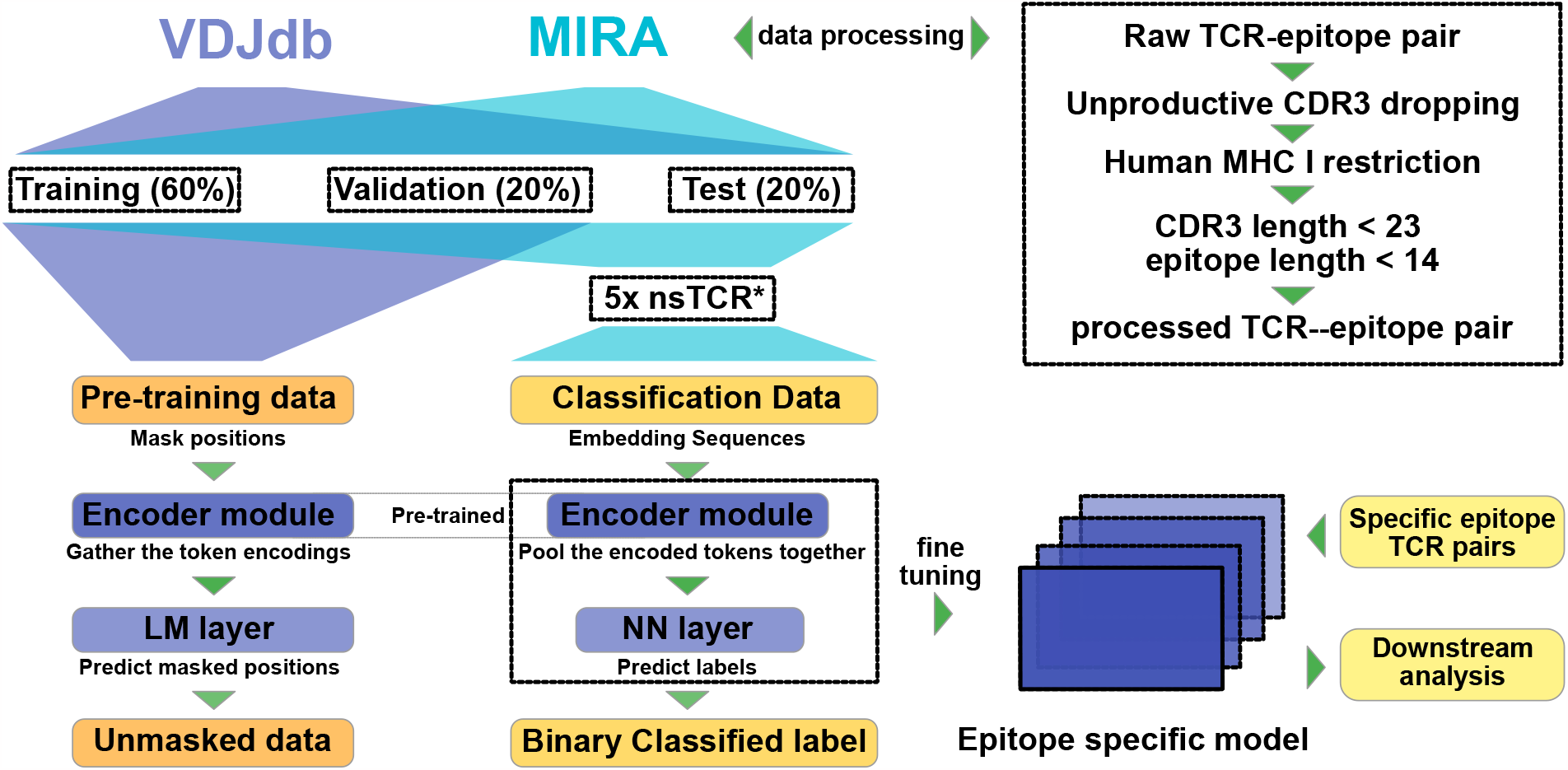
Architecture and performance of the SABRE model. The illustration of the SABRE model.

**Figure 2.**
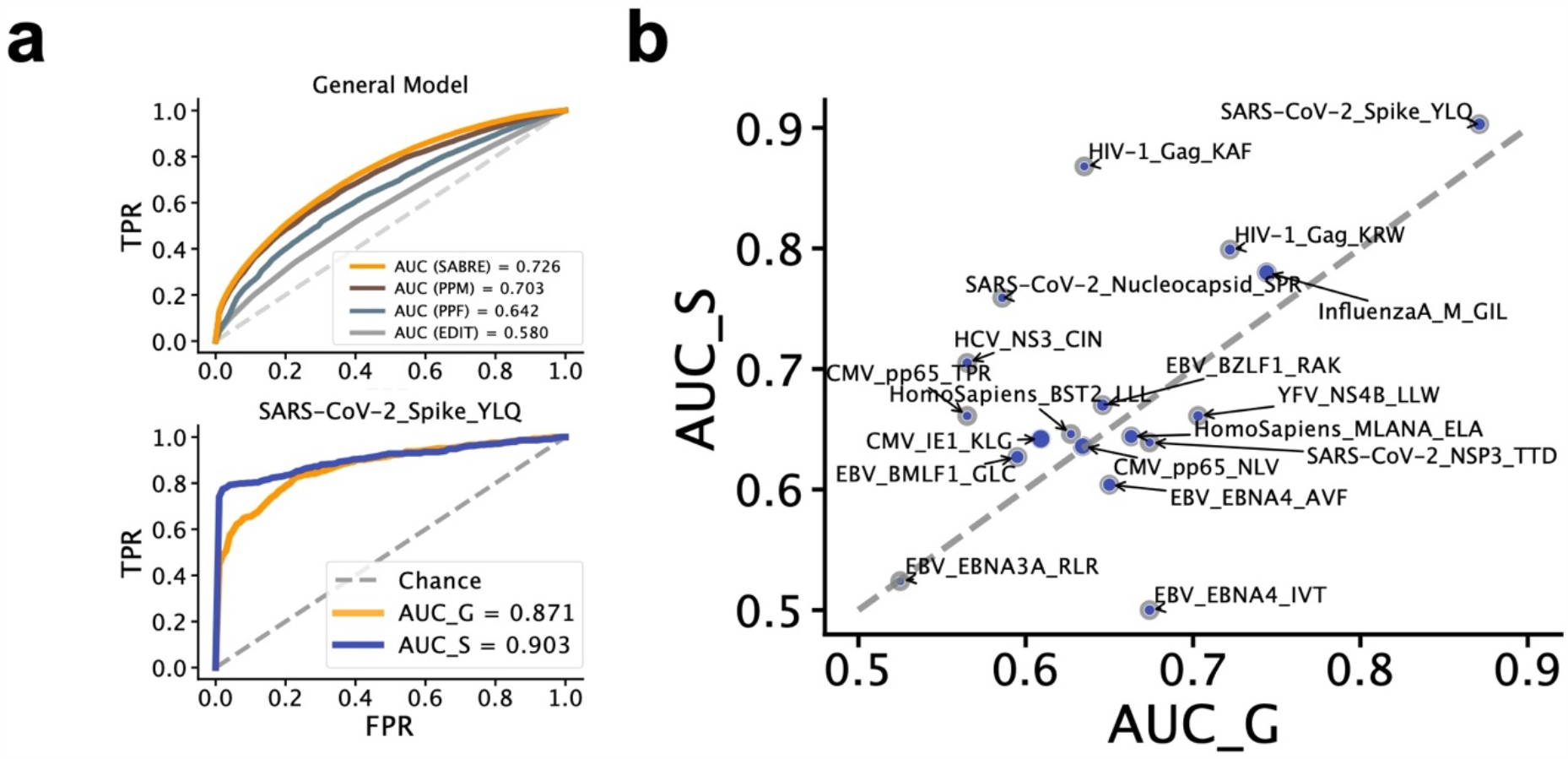
Comparing the general SABRE model with the epitope-specific SABRE model. a. auROC of the SABRE model (SABRE) and SARS-CoV-2 Spike YLQ epitope specific model. Predicting unseen CDR3 compared with the PanPep majority setting (PPM), few-shot setting (PPF), and the edit distance-based method (EITD). *nsTCR, non-specific TCR, which is unknown interaction with any epitope. b. The AUC of 18 high-abundance epitopes’ fine-tuned models compared (AUC_S) with the general prediction model (AUC_G).

**Figure 3.**
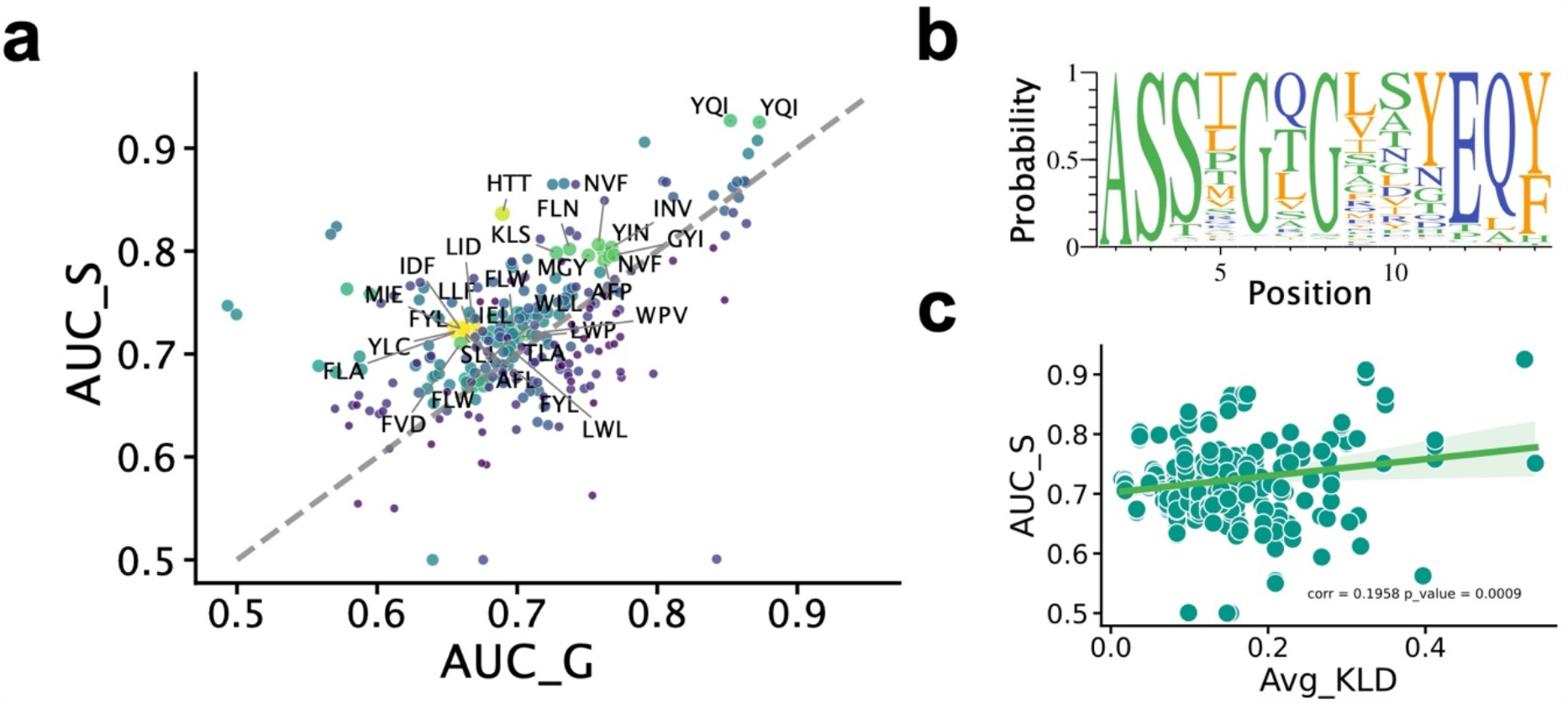
Epitope-specific model application in MIRA dataset. a General model prediction versus epitope-specific model prediction. b. logo plot of YQI-related 15 aa TRB CDR3 sequences. c. Average KL divergence of 15 aa CDR3 sequences starting from 4^th^ position to 10^th^ position of each epitope versus related specific model prediction AUC. The KL divergence is calculated by computing specific epitope-related TRB CDR3 sequences (test distribution) against all TRB CDR3 sequences in the database (background distribution).

The performance of the epitope-specific model depends on the training sample size (number of TCRs) and the specificity of TCR-epitope pairs. AUC increased sharply from 500 to 1,500 TCRs for training and testing the model (training and validation and test ratio are 3:1:1). Further increased sample size would yield diminishing returns (Supplementary Figure 7. Additionally, We found that the YQIGGYTEK-related specific model achieved an AUC of over 0.9, exceeding any other specific model performance at any sampling size level.

To examine the consistency of TRB CDR3 sequences related to the YQI epitope, we observed that the CDR3 sequences in the middle had two meager diversity positions dominated by glycine residues at the 6^th^ and 8^th^ positions (Figure 3b). Apart from the length equal to 15, we also found that in some majority lengths, such as 14, 16, and 17, there were two high-probability glycine residues in the middle of CDR3 sequences. These dominant amino acids make remembering and establishing associations with the specific epitope easier for the language model.

However, when we compared YQI-related CDR3 sequences with all records in the MIRA database, we found that glycine dominance at the 8^th^ position was common. Therefore, we evaluated the entropy of each amino acid position to the experience (across all TRB CDR3 in the database) using KL divergence and found that the YQI-related TRB CDR3s at the 8^th^ KL divergence were smaller than at the 6^th^ position. Moreover, we found that some epitope-related TRB CDRs also had several dominant amino acids at some positions. Although these positions had higher KL divergence, the probability of such dominant amino acids didn’t exceed 50%. In this view, the amino acid absolute dominant is the key to predicting the TCR epitope binding. This means the CDR3 sequences must have one or more unique amino acids compared to the background, and the percentage of that unique amino acid must show significant dominance.

We also aimed to assess the model’s robustness concerning the CDR3 sequence similarity between the training and test datasets. We used the quantile 0.01 edit distance between training and test TCRs to evaluate the conservativeness of sequences in each epitope pool. The Pearson correlation coefficient between the quantile 0.01 edit distance and the AUC of the fine-tuned model was -0.89 (p-value = 4.9*10-21), indicating that higher sequence similarity between the training and test sets leads to better prediction (Supplementary Figure 8).

Our study utilized transformer encoders to capture amino acid features within our model. We examined TRB CDR3 sequences associated with the HLA A2-restricted HIV epitope SL9 to identify activated TRB CDR3 residues corresponding to the SLYNTVATL epitope sequence ^35^. The hidden state values for each residue exhibited variability (Supplementary Fig. 4). We employed Pearson’s correlation coefficient to represent the relationship between each residue in the CDR3 and epitope sequences across all features. Figure 4a highlights regions with high correlation coefficients, suggesting the involvement of specific residues from CDR3 and epitopes. We compared these high correlation coefficient regions (excluding the borders of CDR3 and epitope) with known hotspots for antigen recognition in the TCR-pMHC complex. In the structural model, a residue bond contact is observed between TCR 97V and residue 6V in the SLYNTVATL epitope ^35^, further corroborating the heatmap’s indication of a strong correlation between 6V in the epitope and the CDR3 sequence.

**Figure 4.**
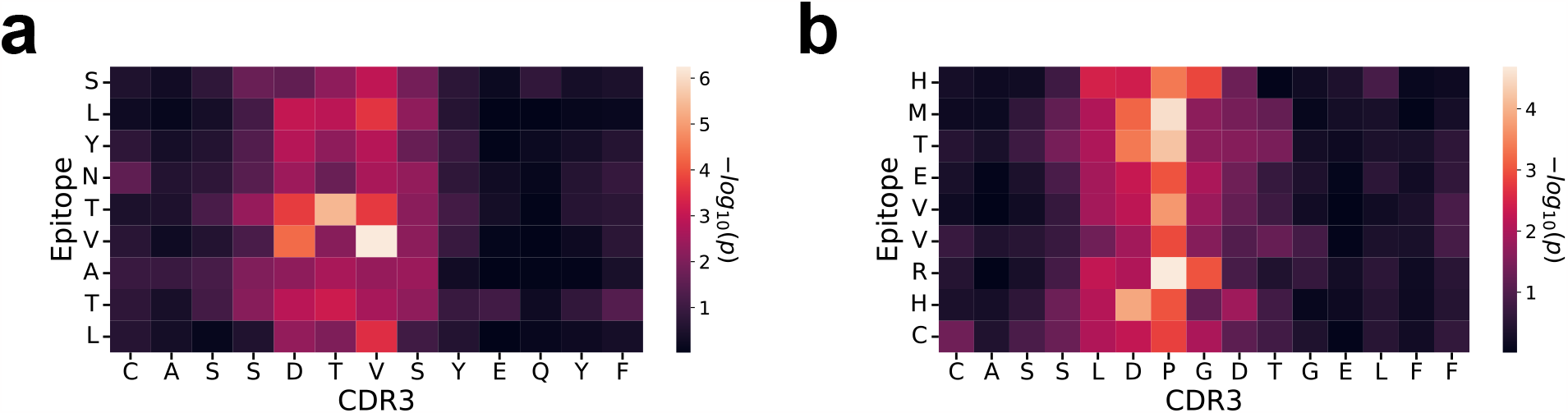
Interpreted the SABRE model intermedial layer output. a-b. The correlation coefficient between CDR3 and epitope’s output of HIV epitope SL9, Neoantigen p53R175H.

Another example involves T cell receptors targeting the p53 cancer neoantigen (HMTEVVRHC). The TCRs specific to p53R175H are diverse, allowing for recognition of the p53 neoantigen and opening potential avenues for developing more effective cancer immunotherapies ^5^. Although the overall number of contacts contributed by CDR3β is smaller than that of CDR3α, we identified that the highest correlated residue pair (P98β and Rp7) from the SABRE transformer encoder output (Figure 4b) accurately reflects the interaction observed in the structural model ^5^. This finding suggests that we can biologically interpret the deep learning output meaningfully or employ a more structured model to enhance prediction accuracy.

While our analysis revealed considerable consistency between the correlation of encoded features for CDR3 and epitope sequences and the interactions demonstrated in the structural model, we cannot definitively conclude that the SABRE model can predict the actual interaction within the structure. Our model may contribute to the task if the sequence reflects the interaction. Additionally, we observed that the same type of residues in the TCR CDR3 and epitope are more likely to exhibit a significant correlation, even though it is not a strict requirement in 3D structure. The interaction between TCRs and epitopes is a complex process influenced by various factors, including the overall shape and charge distribution at the binding interface. Beyond predicting TCR-epitope binding, our method holds the potential for exploring additional capabilities.

The Low-Rank Adaptation (LoRA) technique, pioneered by Microsoft, retains the foundational knowledge of expansive language models by ensuring the conservation of pre-trained weights, thereby minimizing the peril of catastrophic forgetting. Concurrently, it introduces adaptable matrices tailored for domain-specific tasks ^36^. In our research, we integrated LoRA into our original encoder module to enhance the optimization of our epitope-specific model. A comparative analysis between models with and without LoRA revealed that, in general, LoRA did not significantly elevate the Area Under the Curve (AUC) of prediction. However, in instances of suboptimal predictions, the LoRA-enhanced model demonstrated superiority over the archetype. Notably, with epitopes such as RLR and IVT from EBV, the prediction AUC observed an increase from 0.524 to 0.592 and from 0.5 to 0.549, respectively (supplementary figure 9).

The SABRE encoder assimilated sequential, attention-driven, and linkage patterns through its pretraining phase. Subsequently, it underwent a fine-tuning process for the epitope-specific model, modifying the pre-trained model’s weights to hone its task-specific performance. The degree and nature of modifications to the foundational patterns intrinsically depend on the characteristics and volume of the fine-tuning dataset. A heightened diversity in TCR sequences associated with epitopes would invariably introduce complexities to the fine-tuning phase, making precision pattern extraction arduous. Based on the pairwise edit distance of TCRs for specific epitopes, our analysis indicated that the average distance associated with EBV’s RLR-related TCRs surpassed other categories (as illustrated in supplementary figure 10). Given these conditions, it can be postulated that a diminished presence of consensus patterns could render the fine-tuned, epitope-specific model less predictable in its interactions. In such contexts, LoRA offers a pragmatic approach to recalibrating the model, ensuring more accurate prediction outcomes.

The relationship between TCR recognition and HLA presentation is vital for understanding the immune system and developing targeted immunotherapies. The SABRE model currently didn’t consider HLA information, due to a lack of precision HLA type in the MIRA dataset for pretraining. However, as for the pMHC-TCR complete records in the VDJdb dataset, we could retrain the pretraining model and compare how the HLA type could impact the result. Inspired by NetMHCpan, the HLA sequence was encoded using a pseudo-sequence of 34 amino acid residues that come into contact with a peptide ^37^. We extended the model input from 36 to 72 maximum amino acid residues. This way, we could follow the same model architecture and include HLA pseudo-sequences.

In contrast to the epitope-specific model, our analysis revealed that the integration of HLA data could enhance predictive accuracy for specific epitopes, such as KLG/NLV from CMV, RLR/IVT/GLC from EBV, LLW from YFV, and ELA from Homo sapiens (supplementary figure 11). These epitopes, which initially exhibited subpar predictive accuracy in the specific model, displayed notable improvement upon including HLA data, with the prediction AUC for LLW even surpassing a threshold of 0.7. Conversely, adding HLA data did not contribute significantly to performance enhancement for epitopes that already had satisfactory predictions. Integrating HLA information seems to provide a moderating effect, uplifting the lower-tier predictions whilst slightly diminishing the upper-tier ones. A plausible rationale for this phenomenon could be the observed bias in HLA usage (supplementary figure 12). Our examination of the HLA type distribution for each epitope indicated a preponderance of one or two HLA types, which might not offer sufficient diversifying information. Additionally, the uncertainty associated with pinpointing the specific interaction regions between the HLA and TCR means the precise contribution of HLA to predictive improvement remains somewhat nebulous.

The dataset for the MIRA analysis was sourced from various individuals, enabling us to examine the influence of HLA typing on TCR epitope binding. We focused on the singleton epitope HTTDPSFLGRY (HTT) from SARS-CoV-2 ORF1ab, which exhibited the maximum TCR abundance. Within the dataset, TCR interaction records were present for 28 individuals. We narrowed our study sample to 15 individuals with a minimum of 100 TCRs interacting with the HTT epitope.

We encountered an obstacle in identifying the MHC molecule loaded with the epitope, due to the MIRA data’s lack of particular MHC binding information (Supplementary table 2). Consequently, we sought to verify the pMHC complex using the MHC peptide prediction tool, NetMHCpan 4.1^38^. Supplementary Table 1 delineates the number of TCRs, the AUC of each individual’s finetuned model, and the HLA type. Based on previously published research ^39^ and results from the NetMHCpan prediction, we deduced that the HTT epitope was likely loaded on HLA-A01:01. The prediction tool assigned an EL score of 0.925, suggesting a robust binding. Notably, each individual in our study was characterized by the HLA-A01:01 type, which provided a straightforward explanation for the TCR’s ability to interact with HTTDPSFLGRY.

**Table 1.**
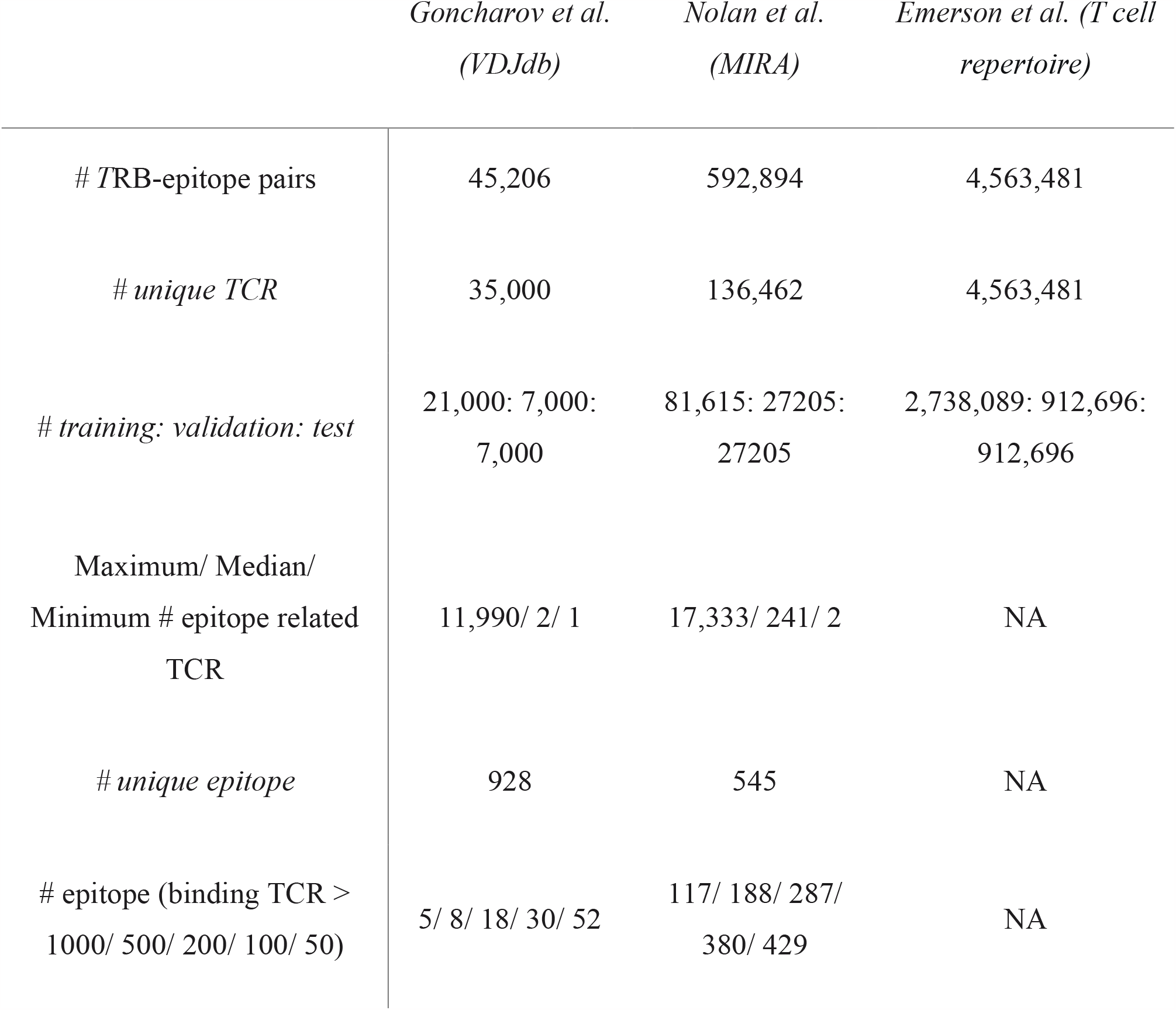
Dataset Details.

Despite the consistency in epitope loading MHC types across all individuals, we observed variability in prediction performances. Other HLA types, such as A*29:02:01 (supplementary table 2), demonstrated weak binding affinity with an EL score of 0.6123. However, this did not necessarily translate into notably high performance, as shown in supplementary Table 1. It was also evident that sample size substantially impacted model performance, with more extensive data sets yielding more reliable results. This observation is corroborated by the Pearson correlation between the log10 ratio of TCR and the AUC, which is 0.786 (p-value = 3e10-4).

Numerous potential factors can influence the accuracy of binding prediction. However, given the known HLA binding type, we postulate that the quantity of training data is of higher importance than differences in HLA type.

## Discussion

The elucidation and prediction of TCR-peptide binding are integral to our comprehensive understanding of human health and disease interventions. As principal components of adaptive immunity, T cells delineate responses against external pathogens and mutated endogenous cells. However, when these T cells misidentify endogenous antigens, they may precipitate autoimmune diseases or hypersensitivity disorders. An accurate predictive model for TCR-peptide binding offers insights into these molecular aberrations, facilitating potential preventive and therapeutic interventions. Furthermore, mastering this predictive capacity can identify novel therapeutic targets and anticipate immune responses to emergent pathogens. Thus, predicting TCR-peptide interactions is not merely foundational in immunology but pivotal for advancing proactive healthcare methodologies.

Identifying T cell epitopes, sections of antigens that T cells or immune system molecules recognize, is key to understanding immune responses to pathogens and allergens. Bioinformatics advancements have enhanced epitope identification, with machine learning techniques predicting TCR-epitope interactions, thus deepening our understanding of the immune response. This is pivotal for creating vaccines, diagnostics, and immunotherapies ^4,19,40,41^. These methods aim to identify the intrinsic relationships within TCRs and distinguish specific TCRs from the background.

In addition to considering the TCR and epitope sequences, methodologies also account for sequences in the V region, including CDR1 and CDR2, which interact with the TCR-pMHC complex. While these regions generally contact the MHC, CDR3 regions mainly interact with the peptide. Although certain CDR1 and CDR2 parts also connect with peptides, many studies use the V gene to represent these parts ^37^. With mass spectrometry improvements, comprehensive datasets of potential T cell epitopes are emerging. Such insights enable a more holistic understanding of TCR-epitope interplay and improve TCR-epitope interaction predictions. The SABRE model considers CDR3 and epitope sequences together to predict TCR-epitope pairs, overcoming previous limitations.

Considering the broader context of TCRs and epitopes, these approaches provide a more comprehensive understanding of the complex interplay between T cell receptor sequences and their respective epitopes, ultimately enhancing the prediction and analysis of TCR-epitope interactions^18^. The SABRE model treated CDR3 and epitope sequences as a combination to predict any TCR-epitope pair, thus removing limitations in prediction design. When we treated epitope as a category and used edit distance to measure CDR3 similarity, we couldn’t deal with TCR-epitope pair with zero knowledge. While Edit distance-based benchmarks exhibited limitations, such as disregarding the biological relevance of predicted TCR-epitope interactions, they offered a valuable benchmark for comparing the performance of various prediction methods and pinpointing areas that warrant further enhancement.

Various epitopes could be defined based on MHC binding probabilities, and numerous TCRs can be sequenced from different tissues. However, when comparing the number of known pairs, we observed that the number of specific TCRs is significantly larger than that of epitopes. In other words, it is common to find numerous TCRs that recognize the same epitope. As of the latest VDJdb version, there are 33,514 CMV records but only 31 unique epitopes. The limited number of epitopes constrained the SABRE model’s ability to learn from the epitope portion of the data. As the number of TCR-epitope pairs increases exponentially, we plan to retrain the model to keep up with the expanding data landscape.

The data used for training our model primarily originate from viral antigens. However, numerous other sources of antigens, such as cancer neoepitopes, represent a significant area of potential T-cell recognition. These alternative sources of antigens could further expand the applicability and utility of the model, as it could be adapted to address a broader range of immunological challenges beyond viral infections ^42,43^. According to structural studies, T-cell receptors (TCRs) recognize neoantigens in two broad categories. The first category involves the recognition of mutated ‘self’ peptides, while the second category entails the recognition of novel ‘non-self’ peptides modified at anchor residues. A mutation that forms an anchor residue generates a new epitope unknown to the immune system, unlike mutated ‘self’ peptides, which differ by only a single amino acid from an existing ‘self’ epitope. Currently, the SABRE model can distinguish TCRs that recognize unique epitopes, assigning identical TCR scores to wild-types (WT) and mutated epitopes based on the general knowledge acquired from different epitopes ^35,44^. TCR recognition of mutated ‘self’ epitopes primarily focuses on the mutation site while discriminating between WT and mutant peptides. Therefore, if the model aims to identify neoantigens, it is essential to incorporate numerous examples of mutant epitopes and their WT counterparts as negative controls.

Furthermore, epitope specificity is manifested in both TCR recognition and HLA presentation. These two processes are crucial in determining the immune system’s response to various antigens. TCRs must effectively recognize specific epitopes, and HLA molecules must present these epitopes to facilitate T cell activation. Understanding the intricate relationship between TCR recognition and HLA presentation is vital for advancing our knowledge of immune system function and developing targeted immunotherapies field ^37,45-48^. The SABRE model does not take into account HLA information for several reasons. Firstly, the contribution of HLA to the model is lower than that of CDR3 and the V gene. Secondly, some TCR-epitope pairs lack HLA-type information. Finally, HLA molecules mainly interact with CDR1 and CDR2, which are not the primary focus of the SABRE model. As it stands, the SABRE model performs classification directly from the transformer encoder embeddings. The epitope-specific model has successfully enhanced prediction accuracy for some well-characterized TCR-epitope binding events. In future developments, we aim to predict binding strength explicitly and then translate it into binary output. We require a more normalized binding affinity score to replace the binary critical label to achieve this. Alternatively, transfer learning could be employed to improve the current prediction output. These advancements will lead to a more refined and accurate model for understanding TCR-epitope interactions.

So far, several methods have focused on TCR-epitope binding fields ^19,21,37,47,49,50^. However, interpreting the predicting model is still challenging ^51^. Besides model interpretation, how to use the prediction to help verify biological findings or lead to potential vaccine findings is a concern ^7^. In this paper, we dug into the interlayer of the SABRE model, and the interaction between amino acids from TCR and epitope may reflect the correlation of output values. We found that high correlation coefficient sites may get close to the structure level. Another application of the SABRE model is to qualify how the mutations in the epitope could impact the binding. Many variants were generated for the SARS-COVID pandemic outbreak as the virus spreads ^52,53^. Many mutations affected the immune response level for the most recent Omicron variant ^46,54^. We compared the predicted change when known TCR binds to the original or mutated epitope. Those findings may help understand the virus mutation from the TCR interaction aspect ^32,55,56^.

## Method

### Data collection and processing

We assembled a dataset comprising pairs of T-cell receptor (TCR) CDR3 sequences and their associated target epitope sequences, obtained from the VDJdb database on April 4^th^, 2022. To ensure relevance to human immunology, we excluded sequence pairs derived from non-human hosts and those associated with major histocompatibility complex (MHC) class II. The length of CDR3 usually does not exceed 21 amino acids (AAs); the epitope is less than 14 AAs ^31,57^. Subsequently, we concatenated the CDR3 and epitope sequences, limiting the combined length to be less than or equal to 35 amino acids (35,000 out of 35,032 in VDJdb were kept). This limitation was based on the observation that CDR3 sequences typically range between 9 and 22 amino acids, while epitope sequences range between 8 and 13 amino acids. For negative control TCRs, we selected TCRs from non-epitope-specific TCR repertoires ^3^. We also obtained SARS-CoV-2-specific TCR and the antigens from the MIRA Open Access database ^32^ and processed them similarly to VDJdb sequences (583,449 out of 592,894 in MIRA were kept). Data used in this work is located at https://immunerace.adaptivebiotech.com/step-one-understand-sars-cov-2/ https://enrollimmunerace.adaptivebiotech.com/restricted/MIRA-Release001.zip

### Model architecture

We used the labeled data containing the known TCR and epitope binding information for specific binding tasks by fine-tuning the pre-trained transformer model. Due to the benefit of the transformer, the model can handle long-range dependencies between input and output without considering their relative position in the sequence, which can learn and incorporate TCR and epitope as a whole picture. The transformer model encoded the CDR3 and epitope-paired sequences, incorporating padding and masking techniques. We compiled a large dataset comprising known TCR-epitope pairs to pre-train a TCR-epitope transformer encoder. To mitigate potential sequence order bias, as TCR and epitope do not interact in a parallel linear manner in their 3D structure, we employed various combinations: standard order, reversed epitope sequence with standard CDR3 sequence, standard epitope with reversed CDR3 sequence, and both sequences reversed. The sequences were then tokenized by amino acid. Randomly selected tokens were used to establish a “masked language model” task, with each token (excluding epitope/CDR3 separator tokens and padding tokens) having an equal likelihood of being masked. Five positions in the input sequence were masked, and the transformer encoder attempted to predict the masked tokens based on the contextual information from the remaining sequence. The TCR-epitope-specific prediction was treated as a binary classification task. If TCR-epitope binding originated from verified TCR-peptide pairs, the pair was assigned a label of 1; otherwise, a label of 0 was assigned to random shuffling TCR-epitope pairs. The encoded tokens were pooled for the prediction task, and dense layers were added to the transformer encoder. The prediction model was fine-tuned with a lower learning rate (1e-4) than the pretraining step (1e-3). The core of the SABRE model was built using Keras^58^. Keras’s Natural Language Processing (NLP) API was also used for the transformer block setup. The transformer encoder used sparse categorical cross-entropy, and the binary classification used binary cross-entropy as a loss function. The transformer encoder has three blocks and four heads in each block. We set 50 epochs in the pretraining step and 25 epochs in the finetuning step.

### Benchmark method and performance metrics

Numerous benchmark methods have been proposed to assess the performance of TCR-epitope prediction algorithms, including those founded on sequence distance. True positive rate (TPR), false positive rate (FPR), and accuracy (ACC) are derived from the counts of true positives (TP), true negatives (TN), false positives (FP), and false negatives (FN). The area under the receiver operating characteristic curve (AUROC) is a metric to evaluate our classification model’s performance, gauging the model’s capacity to differentiate between positive and negative classes.

Edit distance, which quantifies the minimum number of operations necessary to transform one character string into another, has been employed as a benchmark to assess the accuracy of TCR-epitope prediction algorithms. Using a predetermined edit distance cutoff, we established a confusion matrix by comparing the training and test datasets. If the distance between the test and training CDR3 sequences equaled the threshold and both CDR3 sequences bound to an identical epitope, we incremented the true positive count. A false negative count was defined when the distance exceeded the threshold, but the two CDR3 sequences belonged to the same epitope category. Consequently, we were able to compute TPR and FPR using analogous definitions.

As an alternative approach to generating a smooth AUROC curve, we transformed the edit distance using the exponential negative edit distance, e^−*d*^, where d represents the edit distance between two CDR3 sequences. For benchmarking the performance of the SABRE prediction model, we assigned a true label value of one if two CDR3 sequences interacted with the same epitope; otherwise, the true label value was set to zero. Furthermore, we expanded the edit distance evaluation to incorporate the k-nearest neighbors (KNN) based prediction method. By selecting an odd number of nearest neighbors, we employed the distance measurement provided by TITAN’s baseline 32 to compute the AUROC ^26^.

The SABRE is also available at https://github.com/ShenLab/SABRE.

## Notes

https://github.com/ShenLab/SABRE

### Competing Interest Statement

The authors have declared no competing interest.

