## supplemental Files for "SABRE: Self-Attention Based model for predicting T-cell Receptor Epitope Specificity"

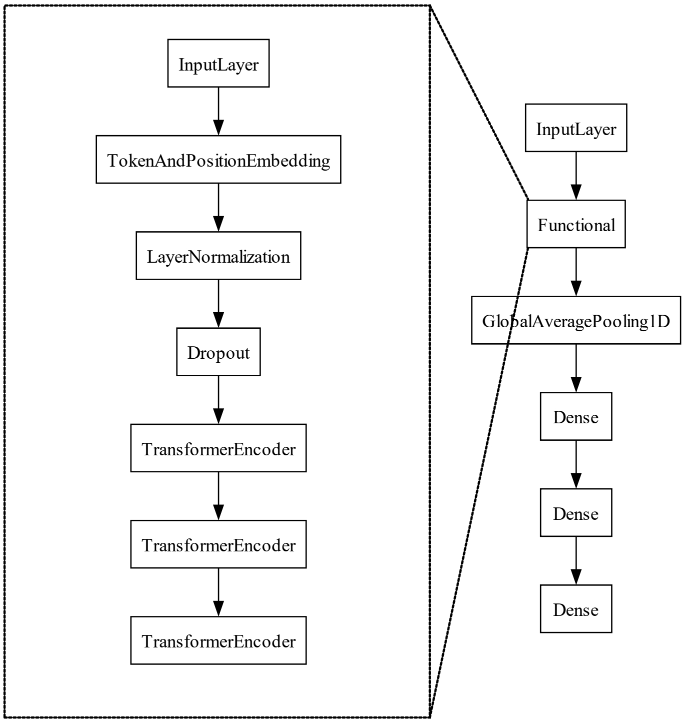


Supplementary Fig. 1 the SABRE model architecture.

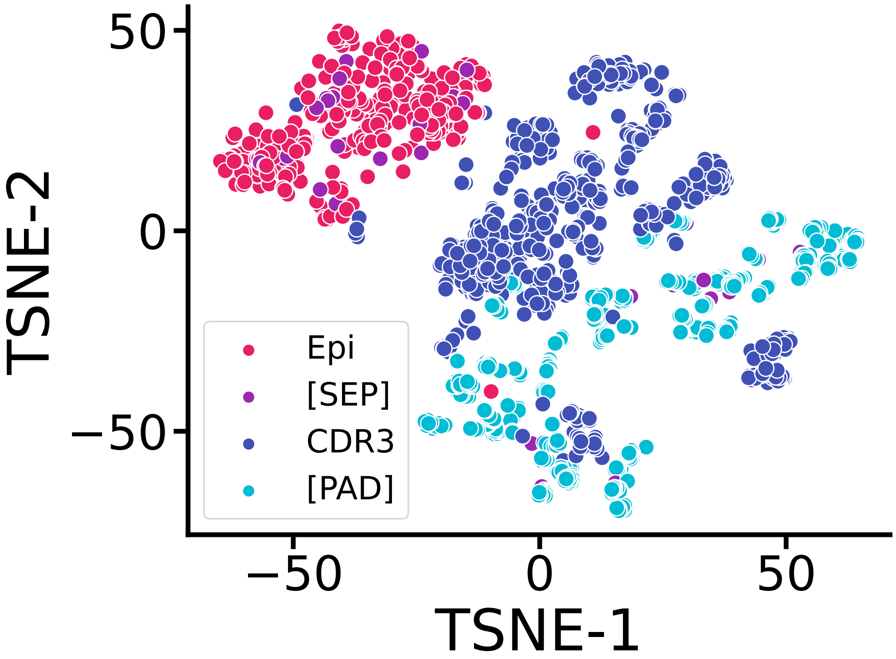


Supplementary Fig. 2 t-SNE plot of transformer encoder embedded epitope and CDR3 sequences.

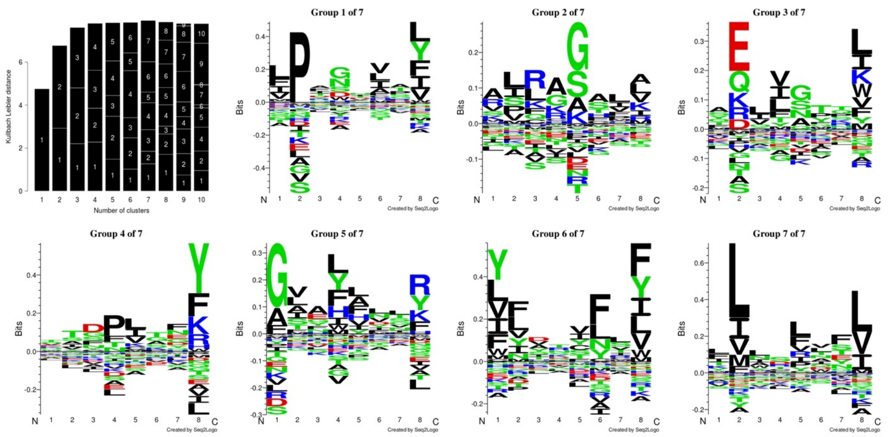

Supplementary Figure 3. Displayed the sequence alignment and motif characterizing seven groups generated with Seq2Logo.

Commented [MOU1]:

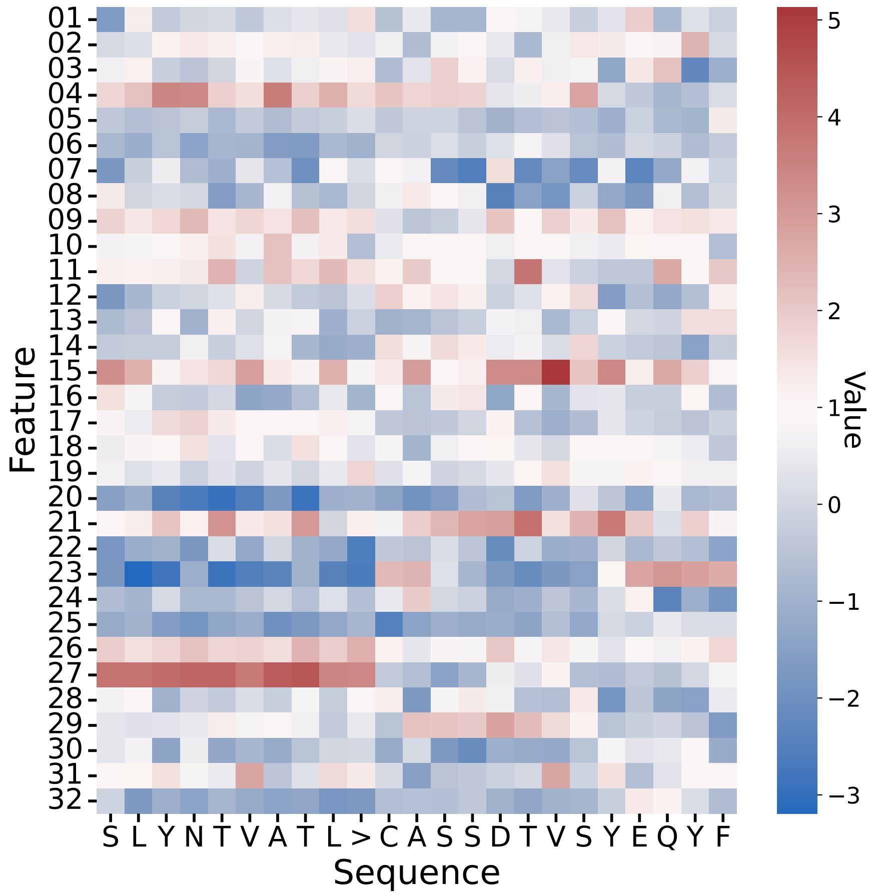

Feature
01 02 03 04 05 06 07 08 09 10 11 12 13 14 15 16 17 18 19 20 21 22 23 24 25 26 27 28 29 30 31 32
S L Y N T V A T L > C A S S D T V S Y E Q Y F
Sequence
Value
5 4 3 2 1 0 −1 −2 −3

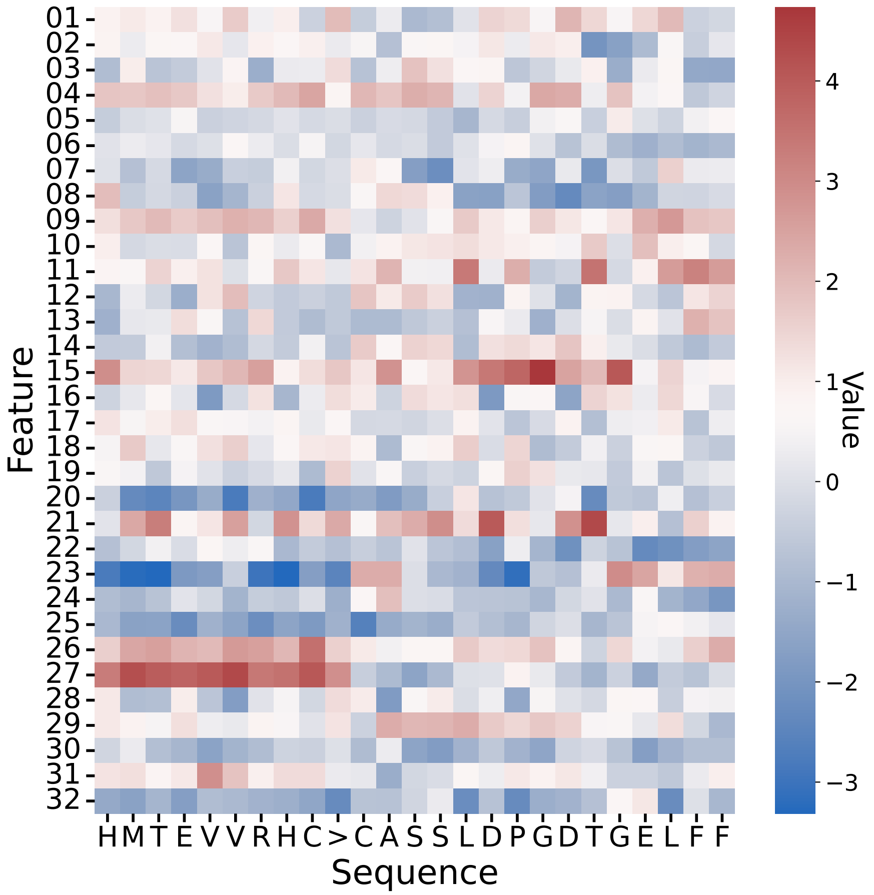


Supplementary Figure 4. Multiple transformer encoder channels' output of HIV epitope SL9, Neoantigen p53R175H, and their corresponding TCR

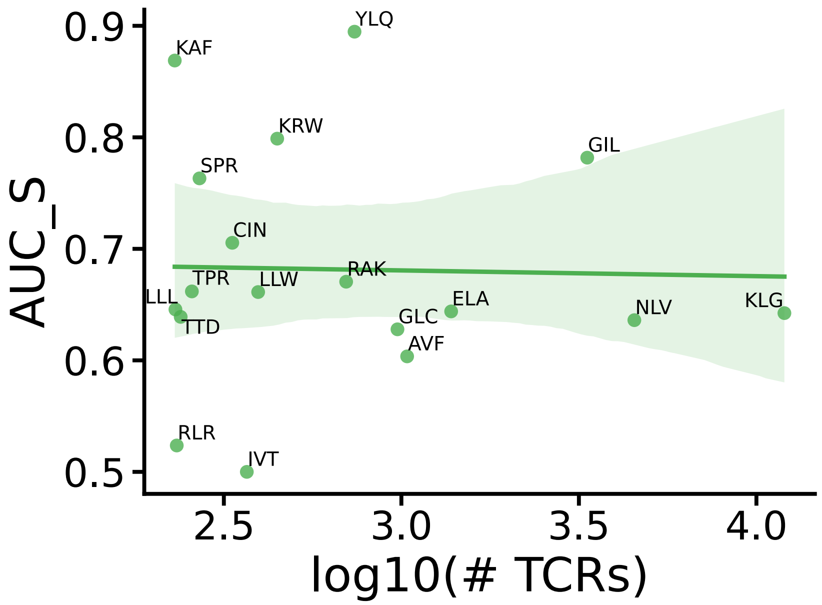


Supplementary Figure 5. Correlation between the number of TCR and AUC for epitope-specific prediction

Using Clustal Omega for multiple TCR CDR3 sequence alignments, we observed that IVT-related TCRs had higher amino acid diversity in the second half than well-predicted epitopes like YLQ (Supplementary Figure 6). Performance variations also arose from HLA-type variation and differences in sequencing methods or quality. We planned to apply the model to the MIRA dataset for subsequent analyses to minimize experimental bias.

**Commented [YS2]:** Does not belong to Results section

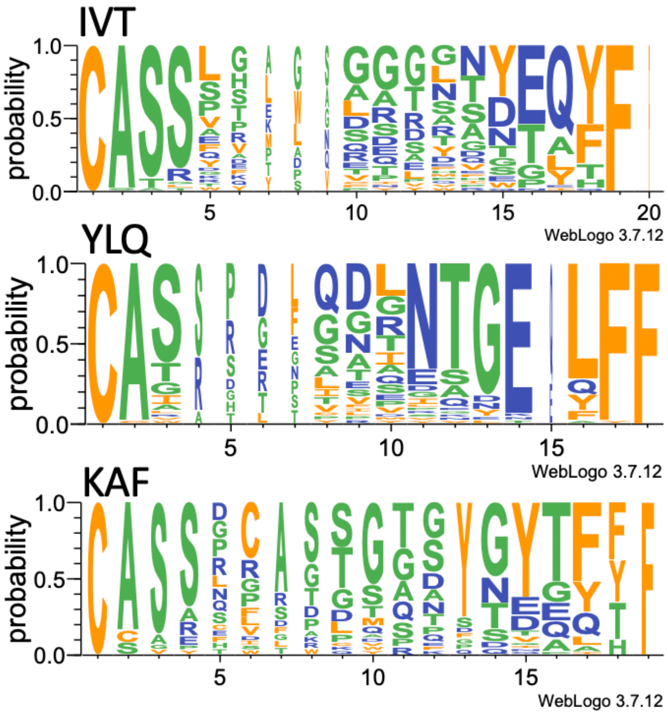


Supplementary Figure 6. Logo plot of three epitope-related TCRs. IVTDFSVIK (IVT), YLQPRTFLL(YLQ), KAFSPEVIPMF(KAF).

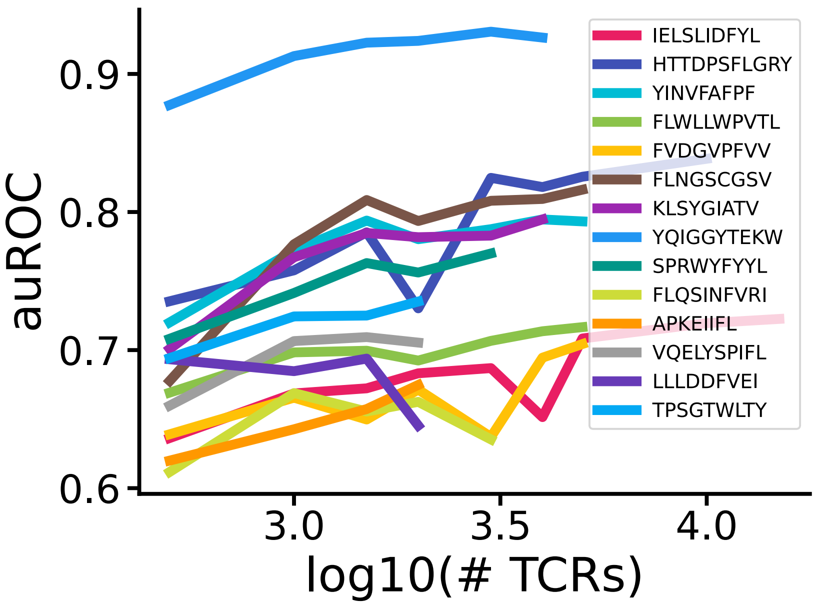


Supplementary Figure 7. Number of TCRs used for training and testing with the correlated finetuned model performance of each high abundance epitope (at least 2000 TCR in MIRA dataset)

(Unseen epitope prediction)

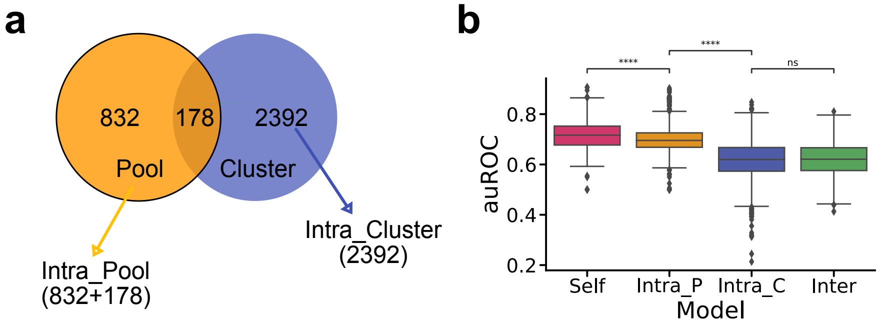

Supplementary Figure 8. Unseen epitope testing. a. two ways for pairwise epitope grouping. b. test performance of each epitope pair.

Our analysis showed that our SABRE model could reasonably predict TCR-epitope pairs when the epitope already exists in the training dataset. In addition, the epitope-specific retrained model further improves prediction power. However, we also found that our model's prediction power drops significantly when the epitope and TCR are from different datasets, especially for training on VDJdb and testing single virus-specific datasets such as the SARS-CoV-2 MIRA dataset.

Due to the unique TCR-epitope identification process, TCRs and epitopes were highly concentrated in subgroups within the MIRA dataset. Therefore, we tested the epitope-specific pairs using several models, including the Self Model (training and test TCRs are related to the same epitope), Intra_Pool Model (training and test TCRs are connected to different epitopes that interact two the same TCRs), Intra_Cluster Model (training and test TCRs are related to different epitopes that are similar sequences and didn’t interact to the same TCRs) and Inter Model (training and test TCRs had no related epitopes). Our test results showed that the difference between the Self and Intra Pool Models was significantly more significant than the Inter Model.

TCR-epitope interaction specificity restricted the number of possible epitopes, but conserved sequence motifs in epitopes ensured high receptor specificity toward their cognate ligands. To investigate how conserved and information-rich amino acid patterns in epitopes may impact TCR binding specificity, we employed the unsupervised epitope motif discovery tool GibbsCluster 2.0 [1] to group epitopes and tested different retraining data split methods to evaluate our SABRE performance.

We clustered Covid peptides into meaningful groups using GibbsCluster 2.0 and set a larger number of clusters to identify more specific clusters. Ultimately, we obtained 48 non-void epitope clusters. This unsupervised clustering method had some overlaps with epitopes in the same pool. After excluding the two epitopes from the same pool, we tested one epitope-related pair using the same cluster epitope-specific model. The results showed no significant difference compared to testing with the Inter Model, indicating that epitope sequence similarity does not lead to reliable prediction.

This analysis highlights the challenges of predicting TCR-epitope interactions when dealing with different datasets. Furthermore, it emphasized the need for further research to improve the accuracy and generalizability of prediction models like our SABRE.

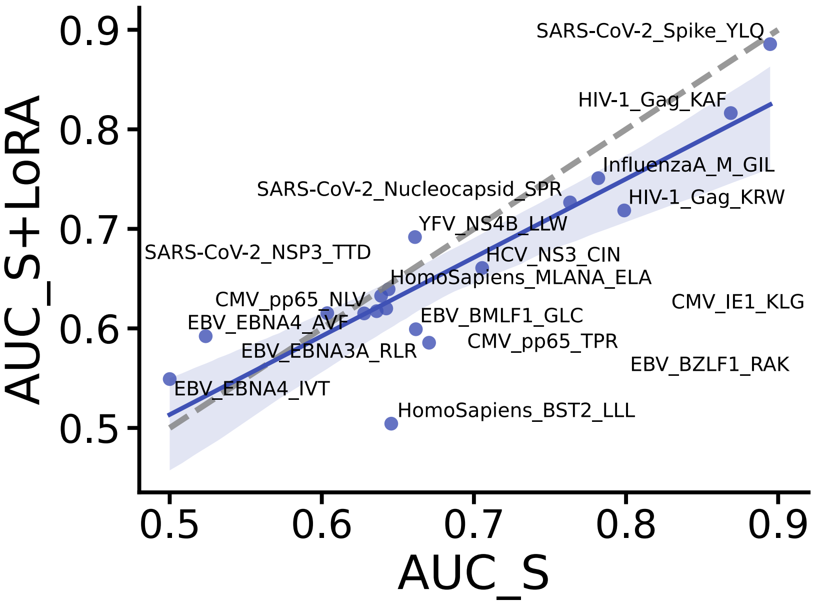


Supplementary figure 9. Correlation between the AUC of epitope-specific prediction and the AUC of specific model with LoRA.

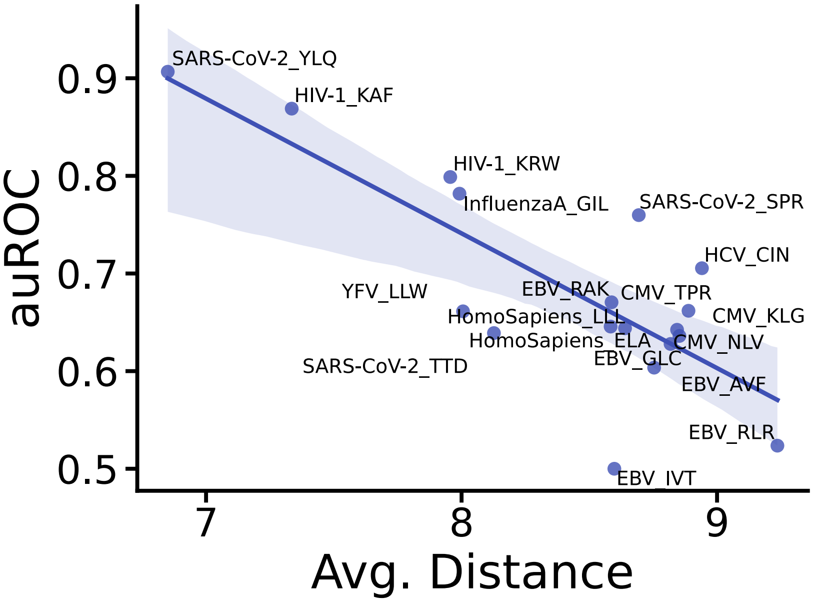


Supplementary figure 10. Correlation between average edit distance of TCRs and AUC for epitope-specific prediction

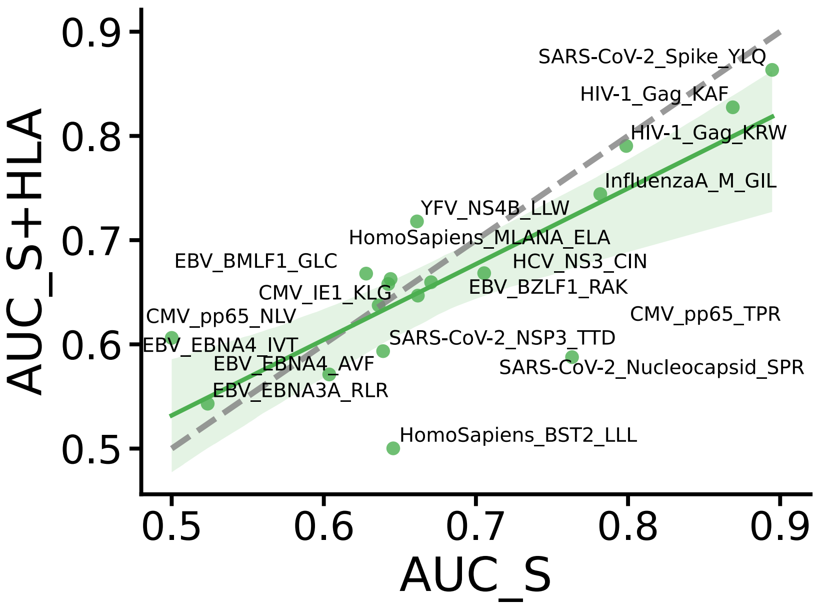


Supplementary figure 11. Correlation between the AUC of epitope-specific prediction and the AUC of specific model with HLA information.

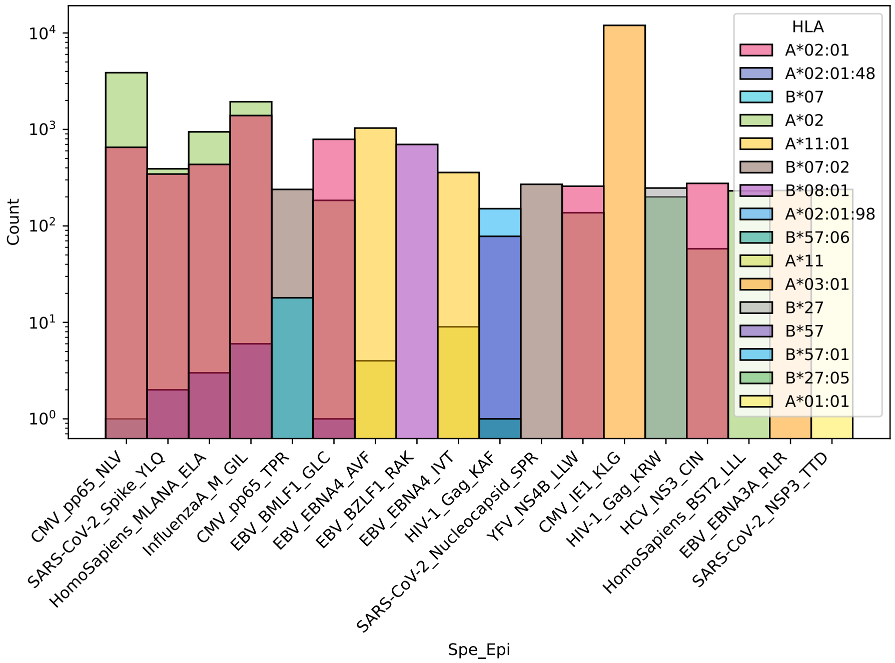


Supplementary figure 12. HLA type distribution of each test epitope from VDJdb.

Supplementary table 1

| | | | EL score | | | | | |
|---|---|---|---|---|---|---|---|---|
| ***Exp.*** | ***#TCR*** | ***AUC*** | **HLA-A** | **HLA-A.1** | **HLA-B** | **HLA-B.1** | **HLA-C** | **HLA-C.1** |
| *eQD111* | 2972 | 0.786 | 0.9251 | 0.0673 | 0.0001 | 0.0002 | 0.0034 | 0.0012 |
| *eMR13* | 1146 | 0.801 | 0.9251 | 0.0003 | 0.0002 | 0.0001 | 0.0003 | 0.0034 |
| *eQD125* | 973 | 0.816 | 0.9251 | 0.0673 | 0.1140 | 0.0006 | 0.0003 | 0.0003 |
| *eMR17* | 875 | 0.805 | 0.9251 | 0.6123 | 0.0003 | 0.3402 | 0.0003 | 0.0019 |
| *eQD114* | 829 | 0.788 | 0.9251 | 0.0003 | 0.0002 | 0.0001 | 0.0034 | 0.0036 |
| *eMR12* | 788 | 0.786 | 0.9251 | 0.0004 | 0.0001 | 0.0013 | 0.0003 | 0.0274 |

| | | | | | | | | |
|---|---|---|---|---|---|---|---|---|
| *eQD121* | 776 | 0.783 | 0.9251 | 0.0003 | 0.0025 | 0.3402 | 0.0019 | 0.0034 |
| *eJL161* | 682 | 0.795 | 0.9251 | 0.0004 | 0.0002 | 0.0018 | 0.0019 | 0.0034 |
| *eQD110* | 523 | 0.787 | 0.9251 | 0.0673 | 0.0032 | 0.3402 | 0.0019 | 0.0019 |
| *eHO134* | 475 | 0.756 | 0.9251 | 0.0003 | 0.0008 | 0.3402 | 0.0019 | 0.0034 |
| *eQD126* | 216 | 0.804 | 0.9251 | 0.0151 | 0.0001 | 0.0002 | 0.0034 | 0.0012 |
| *eQD137* | 186 | 0.657 | 0.9251 | 0.0004 | 0.0020 | 0.0038 | 0.0019 | 0.0274 |
| *eLH47* | 175 | 0.700 | 0.9251 | 0.0004 | 0.0001 | 0.0002 | 0.0034 | 0.0012 |
| *eQD120* | 172 | 0.780 | 0.9251 | 0.0185 | 0.0002 | 0.0001 | 0.0003 | 0.0034 |
| *eHO128* | 142 | 0.693 | 0.9251 | 0.0004 | 0.0002 | 0.0032 | 0.0019 | 0.0034 |
| *eJL162* | 129 | 0.694 | 0.9251 | 0.0004 | 0.0001 | 0.0006 | 0.0003 | 0.0003 |

Supplementary table 2

| ***Exp.*** | ***#TCR*** | ***HLA-A*** | ***HLA-A.1*** | ***HLA-B*** | ***HLA-B.1*** | ***HLA-C*** | ***HLA-C.1*** |
|---|---|---|---|---|---|---|---|
| *eQD111* | 2972 | A*01:01:01 | A*11:01:01 | B*07:02:01 | B*08:01:01 | C*07:01:01 | C*07:02:01 |
| *eMR13* | 1146 | A*01:01:01 | A*24:02:01 | B*08:01:01 | B*40:01:02 | C*03:04:01 | C*07:01:01 |
| *eQD125* | 973 | A*01:01:01 | A*11:01:01 | B*15:02:01 | B*55:02:01 | C*01:02:01 | C*08:01:01 |
| *eMR17* | 875 | A*01:01:01 | A*29:02:01 | B*56:01:01 | B*57:01:01 | C*01:02:01 | C*06:02:01 |
| *eQD114* | 829 | A*01:01:01 | A*24:02:01 | B*08:01:01 | B*41:01:01 | C*07:01:01 | C*17:01:01 |
| *eMR12* | 788 | A*01:01:01 | A*02:01:01 | B*40:01:02 | B*52:01:02 | C*03:04:01 | C*16:01:01 |
| *eQD121* | 776 | A*01:01:01 | A*24:02:01 | B*18:01:01 | B*57:01:01 | C*05:01:01 | C*07:01:01 |
| *eJL161* | 682 | A*01:01:01 | A*02:01:01 | B*08:01:01 | B*13:02:01 | C*06:02:01 | C*07:01:01 |
| *eQD110* | 523 | A*01:01:01 | A*11:01:01 | B*44:02:01 | B*57:01:01 | C*05:01:01 | C*06:02:01 |
| *eHO134* | 475 | A*01:01:01 | A*24:02:01 | B*49:01:01 | B*57:01:01 | C*06:02:01 | C*07:01:01 |
| *eQD126* | 216 | A*01:01:01 | A*03:01:01 | B*07:02:01 | B*08:01:01 | C*07:01:01 | C*07:02:01 |
| *eQD137* | 186 | A*01:01:01 | A*02:01:01 | B*37:01:01 | B*44:03:01 | C*06:02:01 | C*16:01:01 |

| | | | | | | | |
|---|---|---|---|---|---|---|---|
| *eLH47* | 175 | A*01:01:01 | A*02:01:01 | B*07:02:01 | B*08:01:01 | C*07:01:01 | C*07:02:01 |
| *eQD120* | 172 | A*01:01:01 | A*31:01:02 | B*08:01:01 | B*40:01:02 | C*03:04:01 | C*07:01:01 |
| *eHO128* | 142 | A*01:01:01 | A*02:01:01 | B*08:01:01 | B*44:02:01 | C*05:01:01 | C*07:01:01 |
| *eJL162* | 129 | A*01:01:01 | A*02:01:01 | B*40:01:02 | B*55:01:01 | C*03:03:01 | C*03:04:01 |
